# Sarcoplasmic reticulum remodeling through autophagosome biogenesis during *Drosophila* metamorphosis

**DOI:** 10.64898/2026.10.01.755830

**Authors:** Yuri Kasai, Shreya Borthakur, Sabrina Bennani, Yuriko Sakamaki, Tadayoshi Murakawa, Naonobu Fujita

**Affiliations:** School of Life Science and Technology, Institute of Science Tokyo, Yokohama, Japan; Biomedical Center, Institute of Science Tokyo, Tokyo, 113-8510, Japan; Cell Biology Center, Institute of Integrated Research, Institute of Science Tokyo, 4259 S2-11 Nagatsuta, Midori, Yokohama, 226-8501, Japan

**Keywords:** Muscle, *Drosophila*, Remodeling, Sarcoplasmic Reticulum, Autophagy

## Abstract

The sarcoplasmic reticulum (SR) is an elaborate membrane network and the primary intracellular Ca^2+^ store for muscle contraction. SR abundance, morphology, and spatial organization must be remodeled during muscle atrophy and hypertrophy, yet the mechanisms underlying this remodeling remain poorly understood. Autophagy has been implicated in SR homeostasis and may contribute to this process. We previously showed that a subset of *Drosophila* larval muscles remodels into adult dorsal internal oblique muscles through an autophagy-dependent process accompanied by extensive SR reorganization. Here, we show that autophagy is essential for maintaining SR abundance and architecture during muscle remodeling. Blocking autophagosome formation caused the accumulation of large aggregates of SR membranes, whereas blocking autophagosome–lysosome fusion led to the accumulation of autophagosomes with SR transmembrane proteins localized to their membranes. Moreover, disrupting LC3-interacting region (LIR)-dependent cargo recognition did not impair SR remodeling, arguing against a major contribution from selective autophagic degradation of the SR. Instead, our findings suggest that SR membrane components are incorporated into autophagosomal membranes during muscle remodeling. We propose that this membrane flow couples the removal of excess SR membrane to autophagosome biogenesis during atrophy, thereby facilitating large-scale SR reorganization during muscle remodeling.

**Summary statement:** SR membrane components are incorporated into autophagosomal membranes, suggesting that autophagosome biogenesis directly contributes to SR remodeling during *Drosophila* muscle metamorphosis.

## Introduction

The sarcoplasmic reticulum (SR) is a specialized form of the endoplasmic reticulum (ER) that functions as the primary intracellular Ca^2+^ store required for muscle contraction. It forms an elaborate membrane network surrounding each myofibril, with specialized domains dedicated to Ca^2+^ release and reuptake. The junctional SR closely associates with transverse (T)-tubules and mediates rapid Ca^2+^ release through ryanodine receptors (RyRs), whereas the longitudinal SR is enriched in sarco/endoplasmic reticulum Ca^2+^-ATPases (SERCAs), which pump cytosolic Ca^2+^ back into the SR (Rossi et al., 2008). Proper control of SR abundance, morphology, and spatial organization is critical for muscle function. Indeed, abnormalities in SR architecture are associated with muscle disorders, most notably tubular aggregate myopathy, which is characterized by the accumulation of densely packed SR-derived membranes and progressive muscle weakness (Rashed and Selcen, 2025; Schiaffino, 2012). SR-derived aggregates are also observed in non-muscle disorders and aged mouse skeletal muscle (Sánchez-González et al., 2022; Weisleder et al., 2006). Despite the physiological and pathological importance of SR organization, the mechanisms that maintain and remodel SR architecture remain poorly understood.

Autophagy, a major intracellular degradation pathway, plays a role in ER homeostasis. Selective autophagic degradation of the ER, termed ER-phagy, is mediated by receptors that link ER membranes to Atg8/LC3-family proteins on autophagic membranes (Chino and Mizushima, 2023; Mochida and Nakatogawa, 2022). The ER is also a major membrane source for autophagosome biogenesis. Atg2, a bridge-like lipid transfer protein, transfers phospholipids from the ER to growing isolation membranes (Osawa et al., 2022; Reinisch et al., 2025). Ultrastructural studies have revealed extensive tubular membrane connections between the ER and elongating isolation membranes (Hayashi-Nishino et al., 2009; Uemura et al., 2014; Ylä-Anttila et al., 2009). In addition, ER-resident lipids and transmembrane proteins can be delivered to autophagic membranes through COPII vesicles in *S. cerevisiae* (Shima et al., 2019). Thus, autophagy and the ER are intimately interconnected: the ER is both a target of autophagic degradation and a source of membrane components for autophagosome formation. However, whether and how autophagy regulates the highly specialized SR remains largely unexplored. Muscle-specific deletion of core autophagy genes, including *ATG5*, *ATG7*, *ATG14*, *ULK1/ULK2*, and *FIP200*, in mice causes pronounced SR abnormalities and impaired muscle function (Li et al., 2021; Masiero et al., 2009; Raben et al., 2008; Wang et al., 2019). Notably, stacked SR membranes accumulate in autophagy-deficient muscle fibers, suggesting an important role for autophagy in maintaining SR architecture, although the mechanism underlying this phenotype remains unknown.

*Drosophila* dorsal internal oblique muscles (DIOMs) provide a genetically tractable system for dissecting muscle remodeling *in vivo*. During metamorphosis, a subset of larval body wall muscles escapes developmental cell death and remodels into adult abdominal muscles through autophagy-dependent atrophy followed by hypertrophy (Fujita et al., 2017). We previously showed that autophagy-mediated clearance of larval mitochondria is critical for this remodeling process (Taoka et al., 2025). The SR also undergoes extensive reorganization during DIOM remodeling (Fujita et al., 2017), but the mechanisms underlying this reorganization and whether autophagy contributes to it remain unexplored. Here, we show that inhibition of autophagosome formation causes extensive SR stacking in DIOMs, recapitulating a hallmark of autophagy-deficient mammalian muscle. We further find that SR transmembrane proteins localize to autophagosomal membranes and that LIR–LDS (LC3-interacting region–LIR-docking site)-dependent selective autophagy is largely dispensable for SR remodeling. These findings suggest that the SR contributes membrane components to autophagosomes during muscle remodeling, revealing an unexpected link between SR remodeling and autophagosome biogenesis.

## Results

### Loss of autophagy induces SR stacking in DIOMs

Loss of autophagy led to the formation of abnormally large clusters of mCD8-GFP-positive membranes in DIOMs during the early stage of metamorphosis (Fig. 1A and B) (Fujita et al., 2017). At this stage, autophagy actively degrades organelles, including mitochondria, in DIOMs (Taoka et al., 2025). Because T-tubules are disassembled during DIOM remodeling and mCD8- GFP labels T-tubules in steady-state muscle cells, we hypothesized that these membrane clusters represented T-tubule-derived structures. We therefore examined the colocalization of mCD8-GFP with the established T-tubule markers Dlg1 and mCh-PH^PLCδ^ in *FIP200* RNAi DIOMs. Contrary to our prediction, the mCD8-GFP-positive clusters were negative for both T-tubule markers (Fig. 1C).

**Figure 1.**
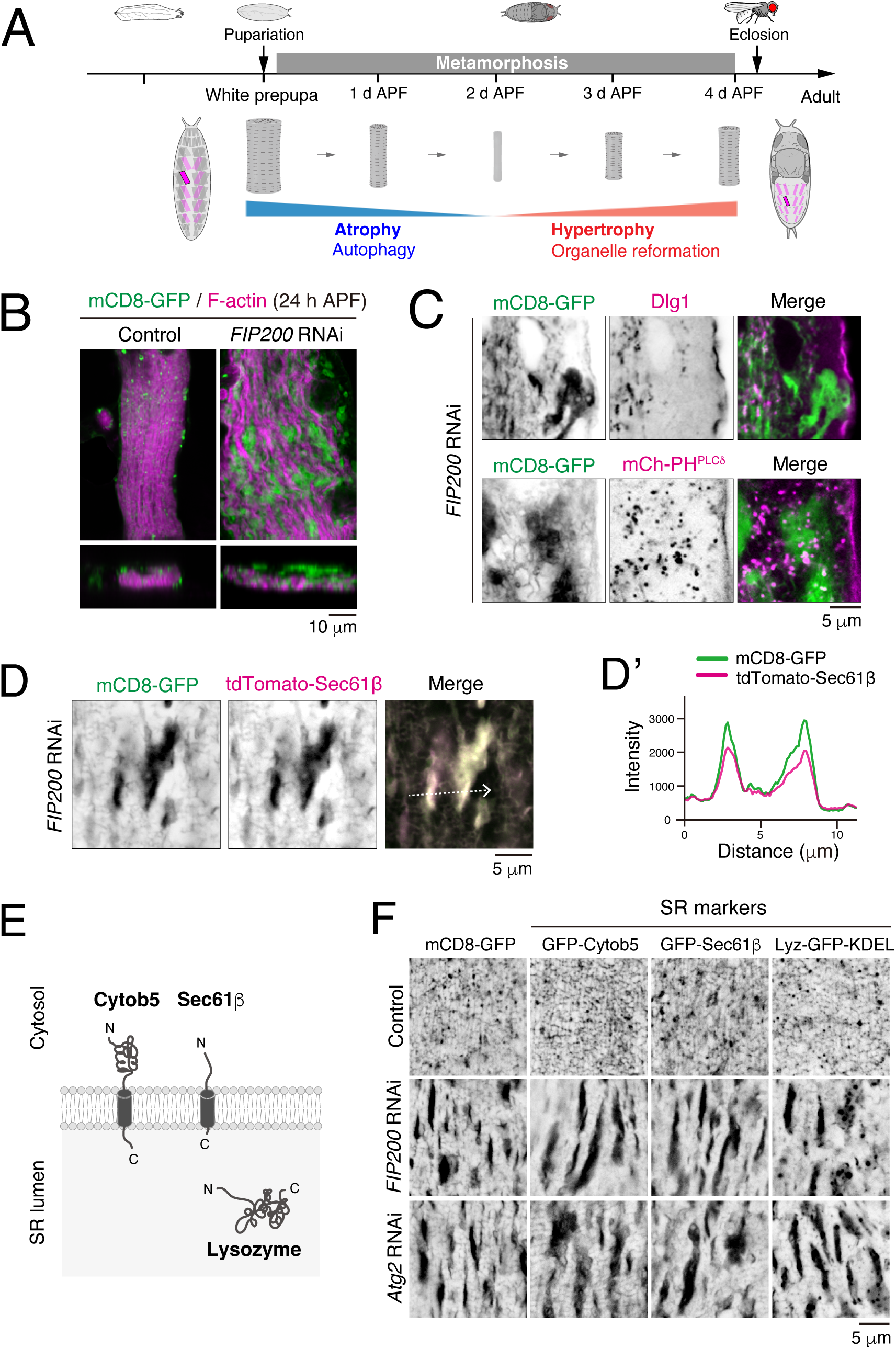
Loss of autophagy induces SR stacking in DIOMs. (A) Timeline of DIOM remodeling during metamorphosis. (B) mCD8-GFP and F-actin in control or *FIP200* RNAi DIOMs at 24 h APF. Projected images of the XY and XZ planes are shown. (C) Colocalization of mCD8-GFP with Dlg1 or mCh-PH^PLCδ^ in 24 h APF DIOMs. (D and D’) Colocalization of mCD8-GFP with tdTomato-Sec61β in DIOMs at 22 h APF. Line profiles along the white dotted arrow are shown. (E) Schematic of the ER/SR markers. (F) Effect of *FIP200* or *Atg2* RNAi on SR markers in DIOMs at 22 h APF.

Although mCD8-GFP is widely used as a plasma membrane marker (Lee and Luo, 1999), it can also label intracellular membranes, including the ER/SR. We therefore examined the colocalization of mCD8-GFP with tdTomato-Sec61β, an ER/SR transmembrane protein. The two markers strongly colocalized within the membrane clusters (Fig. 1D), suggesting that these structures were derived from the SR. To further examine this possibility, we analyzed three GFP-tagged ER/SR markers comprising two transmembrane constructs and one luminal construct (Fig. 1E). Similar to mCD8-GFP, all three ER/SR markers accumulated in stacked membrane structures upon *FIP200* or *Atg2* RNAi (Fig. 1F). Together, these results indicate that loss of autophagy causes abnormal stacking of the SR, rather than T-tubules, in DIOMs.

We examined the dynamics of SR stacking during DIOM remodeling (Fig. 1—figure supplement 1). We observed no apparent difference between control and *Atg2* RNAi third-instar larval body wall muscles. In contrast, GFP-Cytob5-positive aggregates emerged between 13 and 16 h APF in *Atg2* RNAi DIOMs but were not observed in controls (Fig. 1—figure supplement 1A). Live imaging further revealed that these aggregates progressively merged to form larger structures (Fig. 1—figure supplement 1B), suggesting that SR stacks grow by the coalescence of smaller aggregates.

### SR stacking varies depending on the stage of autophagosome formation

ATG proteins function at different stages of autophagosome formation (Fig. 2A) (Nakatogawa, 2020). To determine how disruption of different steps in this process affects SR remodeling, we depleted individual ATG proteins in DIOMs expressing mCD8-GFP and analyzed mCD8-GFP-positive membrane structures at 22 h APF (Fig. 2B). Consistent with the results above, depletion of Atg2, Atg18a, FIP200, Atg1, or Atg9 caused the formation of large mCD8-GFP-positive aggregates. In contrast, the aggregates induced by *Atg5* or *Atg7* RNAi were substantially smaller. To exclude the possibility that the milder phenotypes resulted from inefficient knockdown of Atg5 or Atg7, we compared GFP-Cytob5-positive SR aggregates in *Atg9*-null and *Atg5*-null DIOMs (Fig. 2C). Consistent with the RNAi experiments, SR aggregates were markedly smaller in *Atg5*-null than in *Atg9*-null DIOMs. These results indicate that disruption of early steps in autophagosome biogenesis causes more pronounced SR stacking than disruption of the ubiquitin-like conjugation machinery.

**Figure 2.**
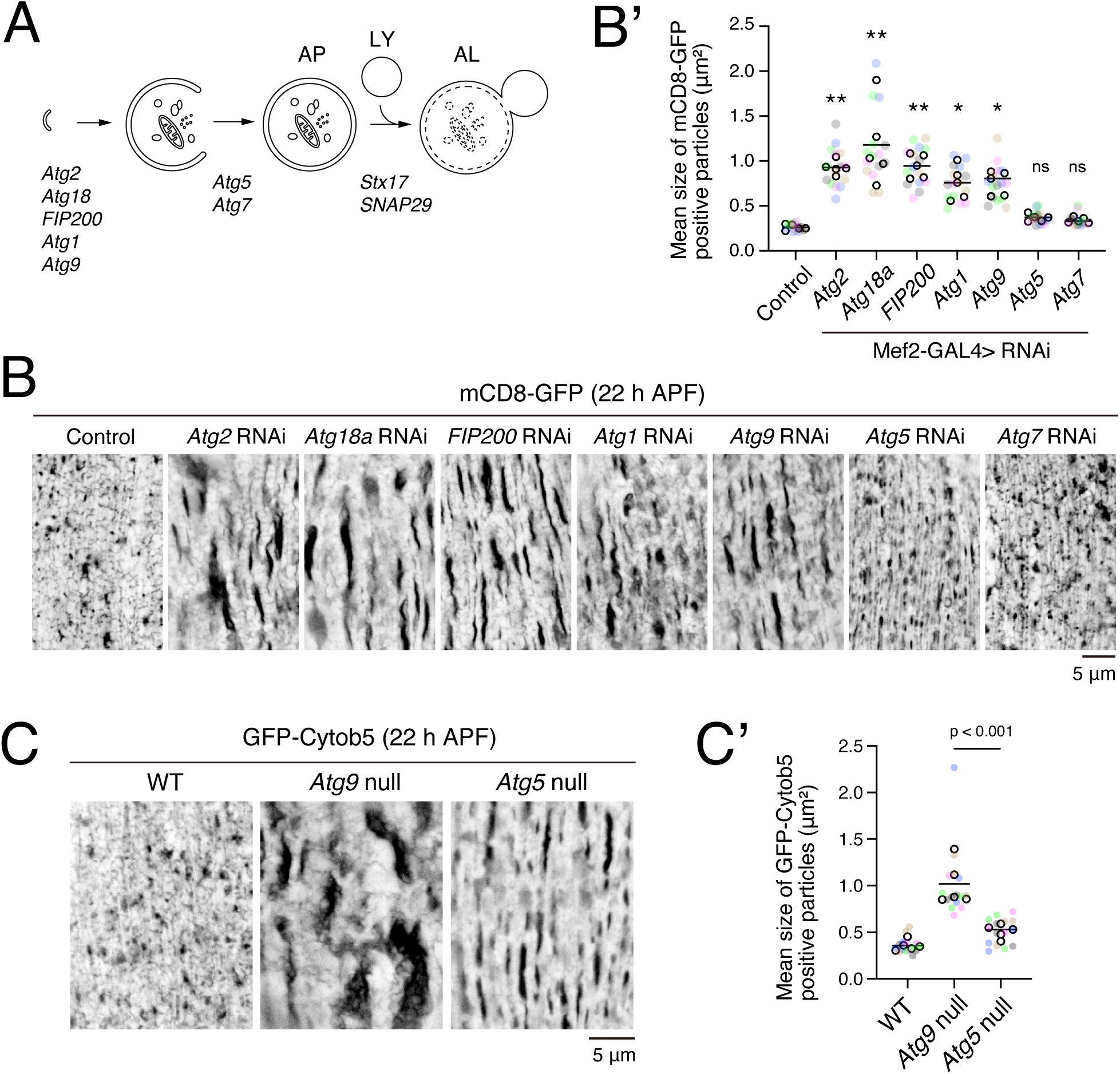
SR stacking varies depending on the stage of autophagosome formation. (A) Schematic of the autophagy process. (B and B’) Effects of RNAi against the indicated *ATG*s on mCD8-GFP-positive structures in DIOMs at 22 h APF. The graph shows the mean size of mCD8-GFP-positive structures. n = 5 animals (15 DIOMs) (Dunnett’s test). ns, not significant; *P < 0.01; **P < 0.0001. (C and C’) GFP-Cytob5-positive structures in *Atg9-* or *Atg5*-null DIOMs at 22 h APF. The graph shows the mean size of GFP-Cytob5-positive structures. n = 5 animals (15 DIOMs) (Dunnett’s test).

We then analyzed SR morphology at the ultrastructural level. Transmission electron microscopy revealed stacked SR membranes in DIOMs lacking ATG proteins at 22 h APF (Fig. 3A–C and Fig. 3—figure supplement 1). Stacked SR membranes were observed in *Atg9*-null, *FIP200*-null, and *Atg5*-null DIOMs but not in wild-type (WT) DIOMs. As previously reported, autophagosomes were observed in WT DIOMs (Fujita et al., 2017). Notably, autophagic membrane structures were also present in *Atg5*-null DIOMs, in which SR stacks were smaller than those in *Atg9*- or *FIP200*-null DIOMs. These observations suggest that the extent of SR stacking is associated with the stage at which autophagosome formation is disrupted.

**Figure 3.**
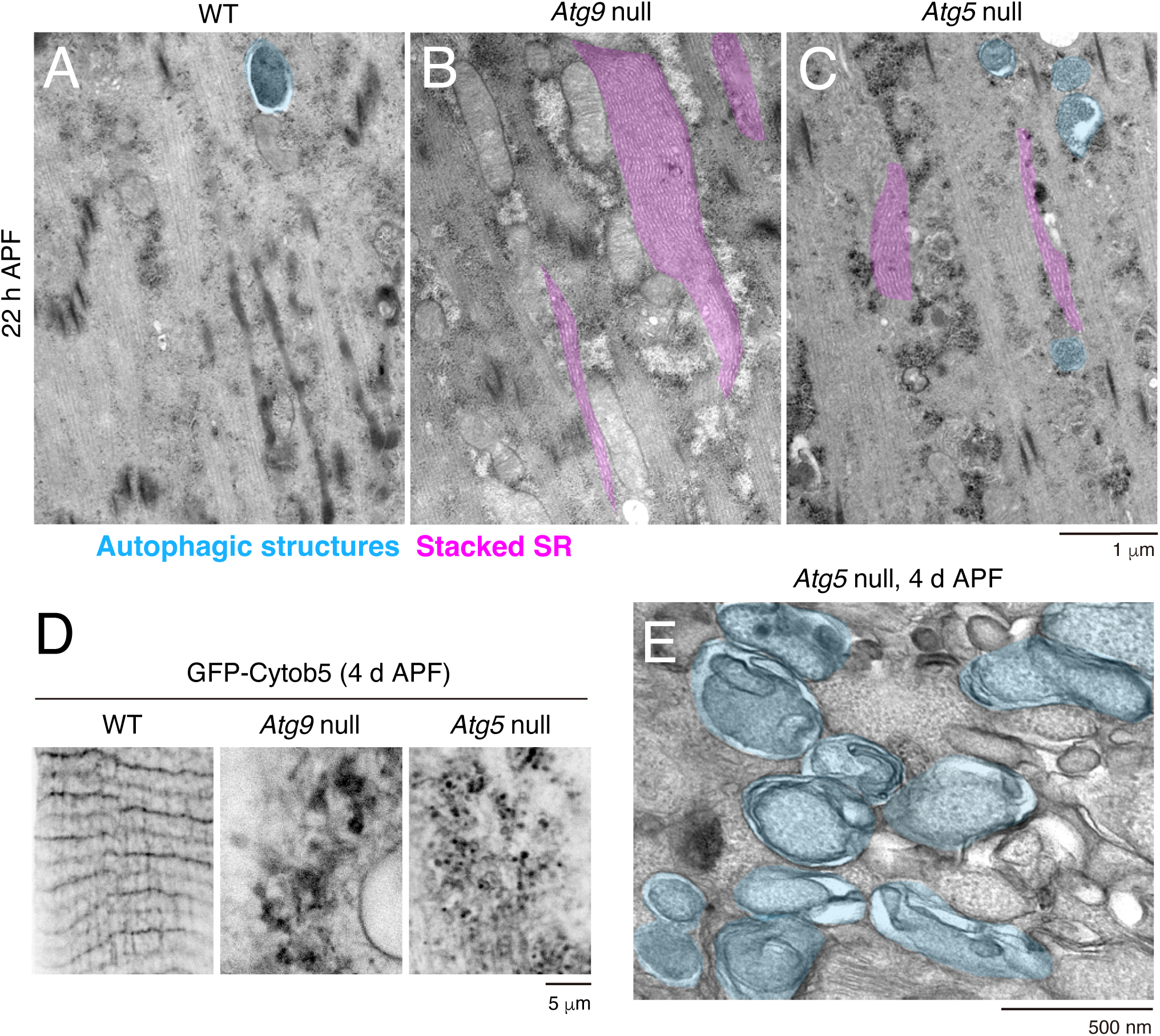
Ultrastructural analysis of *ATG*-null DIOMs. (A-C) TEM images of longitudinal sections of (A) WT, (B) *Atg9*-null, and (C) *Atg5*-null DIOMs at 22 h APF. Stacked SR, pink; autophagic structures, blue. (D) GFP-Cytob5 in *Atg9*- null and *Atg5*-null DIOMs at 4 d APF. (E) TEM image of a transverse section of an *Atg5*-null DIOM at 4 d APF. Autophagic structures are in blue.

We further examined this relationship at 4 d APF, when DIOM remodeling is complete. At this stage, GFP-Cytob5-positive aggregates accumulated in *Atg9*-null DIOMs, whereas smaller vesicular GFP-Cytob5-positive structures accumulated in *Atg5*-null DIOMs (Fig. 3D). TEM revealed a massive accumulation of autophagic membrane structures in *Atg5*- null DIOMs that was not observed in WT DIOMs (Fig. 3E and Fig. 3—figure supplement 1B). Together, these findings suggest that the extent of SR stacking depends on the stage at which autophagosome formation is disrupted and is inversely associated with the formation of autophagic membrane structures.

### Blockade of autophagosome–lysosome fusion causes accumulation of vesicles positive for SR markers

The results above suggested a close relationship between autophagic membrane formation and SR remodeling. We therefore asked how blocking a later step in the autophagy pathway, autophagosome–lysosome fusion, affects SR morphology. Stx17 and SNAP29 are Qa- and Qbc-SNAREs, respectively, that mediate autophagosome–lysosome fusion (Fig. 2A) (Lőrincz and Juhász, 2020). We depleted Stx17 or SNAP29 and examined the distribution of the SR markers GFP-Sec61β, GFP-Cytob5, and Lyz-GFP-KDEL (Fig. 1E). In sharp contrast to *FIP200* RNAi, *Stx17* or *SNAP29* RNAi did not induce stacked SR structures at either 22 h or 4 d APF (Fig. 4A and B). SR markers accumulated in punctate and ring-like structures, particularly at 4 d APF (Fig. 4B).

**Figure 4.**
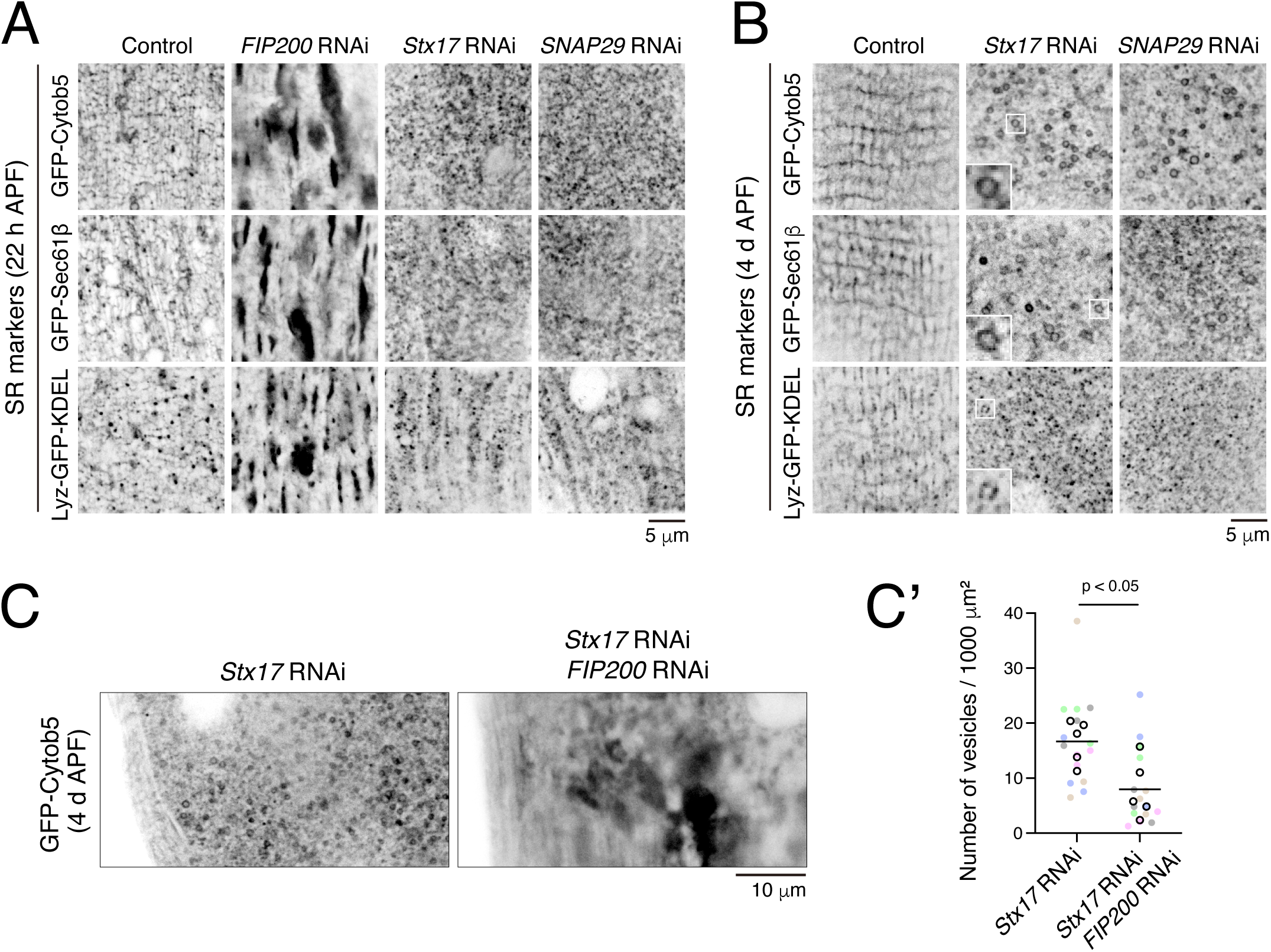
Blockade of autophagosome–lysosome fusion causes accumulation of vesicles positive for SR markers. (A and B) SR markers in DIOMs expressing the indicated RNAi constructs at (A) 22 h APF or (B) 4 d APF. (C and C’) Effect of *FIP200* RNAi on GFP-Cytob5-positive vesicles induced by *Stx17* RNAi in DIOMs at 4 d APF. The graph shows the number of GFP-Cytob5-positive vesicles per unit area. n = 5 animals (15 DIOMs) (Mann-Whitney U test).

We asked whether the formation of these structures positive for SR markers depended on the autophagy machinery. GFP-Cytob5-positive vesicles accumulated in *Stx17* RNAi DIOMs, whereas simultaneous depletion of FIP200 significantly reduced their number (Fig. 4C). Thus, the accumulation of SR marker-positive vesicles upon blockade of autophagosome– lysosome fusion does depend on the autophagy machinery. Together with the SR stacking observed when early autophagosome biogenesis is impaired, these findings raised the possibility that SR membrane components are transferred to autophagic membranes during DIOM remodeling.

### SR transmembrane proteins localize to autophagosomal membranes

To test this possibility, we examined colocalization of tdTomato-Sec61β with autophagosome markers GFP-Atg8a and GFP-Stx17 under conditions in which autophagosome–lysosome fusion was impaired. tdTomato-Sec61β localized to the rims of GFP-Atg8a- and GFP-Stx17- positive autophagic vesicles (Fig. 5A and Fig. 5—figure supplement 1), suggesting that Sec61β localizes to autophagosomal membranes. We next examined this localization at the ultrastructural level by immuno-EM for GFP-Sec61β in *Stx17* RNAi DIOMs. Immunogold labeling for GFP-Sec61β was detected on double-membrane autophagic structures (Fig. 5B), supporting the localization of Sec61β to autophagosomal membranes.

**Figure 5.**
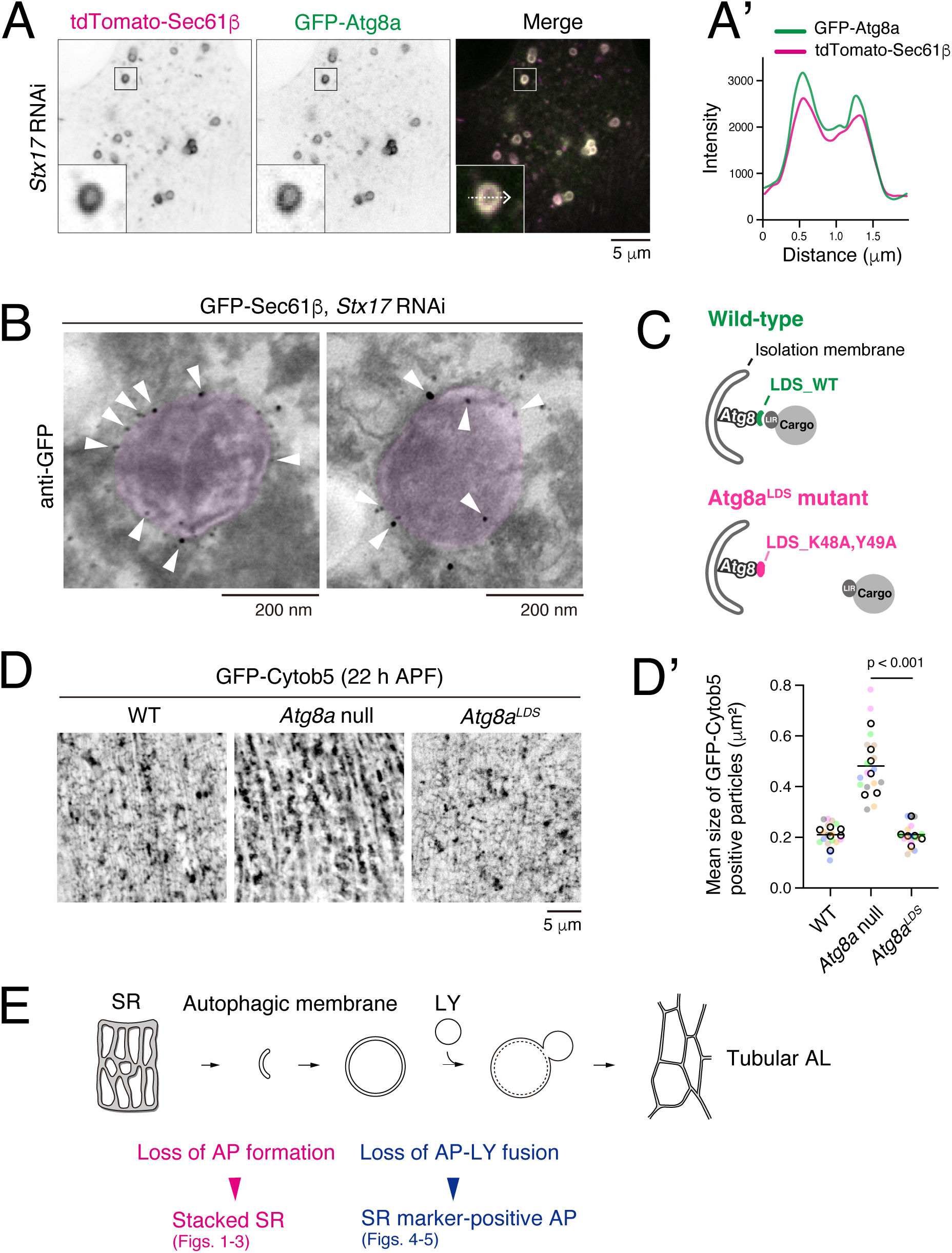
SR transmembrane proteins localize to autophagosomal membranes. (A) Colocalization of tdTomato-Sec61β with GFP-Atg8a in *Stx17* RNAi DIOMs at 4 d APF. Line profiles along the white dotted arrow are shown. (B) Immuno-EM of GFP-Sec61β in *Stx17* RNAi DIOMs. Double-membraned structures are light pink. (C) Schematic of the LIR–LDS- dependent selective autophagy. The Atg8a^LDS^ mutant lacks affinity for LIR motifs. (D and D’) GFP-Cytob5-positive aggregates in *Atg8a*-null and *Atg8a^LDS^* DIOMs at 22 h APF. The graph shows the mean size of GFP-Cytob5-positive structures. n = 6 animals (≥15 DIOMs) (Dunnett’s test). (E) Schematics of autophagy-mediated SR remodeling in DIOMs.

We next examined whether selective autophagic sequestration of the SR contributes to SR remodeling. Known ER-phagy receptors contain LIRs that bind to the LDS of Atg8-family proteins and thereby mediate selective engulfment of the ER (Fig. 5C, top). *Drosophila* has two Atg8 paralogs, Atg8a and Atg8b, of which only Atg8a is expressed in DIOMs (Jipa et al., 2021; Taoka et al., 2025). We therefore examined SR remodeling in the Atg8a^K48A,Y49A^ mutant, where mutations in the LDS disrupt interactions between Atg8a and LIR motifs (Fig. 5C, bottom) (Rahman et al., 2022). If SR stacking resulted primarily from impaired LIR–LDS-dependent selective autophagic degradation of the SR, disruption of this interaction would be expected to cause SR accumulation. Loss of Atg8a resulted in the accumulation of SR stacks at 22 h APF, similar to the phenotype observed in *Atg5*-null DIOMs. In contrast, SR stacking was barely detectable in DIOMs expressing the Atg8a^LDS^ mutant (Fig. 5D). Thus, LIR–LDS-dependent selective autophagy is largely dispensable for SR remodeling in DIOMs. Together with the localization of Sec61β to autophagosomal membranes, these findings support a model in which SR membrane components contribute to autophagosomal membranes during muscle remodeling rather than being sequestered as autophagic cargo through LIR–LDS-dependent selective autophagy.

## Discussion

In this study, we show that autophagy is essential for maintaining SR abundance and architecture in *Drosophila* muscle, consistent with previous observations in mammalian skeletal muscle (Li et al., 2021; Masiero et al., 2009; Raben et al., 2008; Wang et al., 2019). Our findings further suggest an unexpected mechanism by which autophagy contributes to SR remodeling. The SR appears to supply membrane components to autophagosomes rather than being solely sequestered as autophagic cargo. We therefore propose that SR remodeling involves a membrane flow in which SR components are transferred to autophagic membranes and subsequently delivered to lysosomes. Such a mechanism would directly couple the removal of excess SR membrane to autophagosome biogenesis during atrophy, providing an efficient way to coordinate large-scale organelle reorganization during development.

One of the most intriguing observations was the localization of SR transmembrane proteins, including Sec61β and Cytob5, to autophagosomal membranes when autophagosome– lysosome fusion was impaired. Despite the close association of the ER with forming autophagosomes, ER-resident transmembrane proteins in multicellular organisms, including *Drosophila*, are generally thought to be excluded from autophagosomal membranes, which are unusually poor in integral membrane proteins (Baba and Osumi, 1995; Fengsrud et al., 2000). Our data suggest that SR membrane components are incorporated into autophagic membranes during muscle remodeling. How are SR transmembrane proteins transferred to autophagic membranes? One possibility is vesicular transport. COPII vesicles, best known for mediating ER-to-Golgi transport, have also been implicated in autophagosome biogenesis. In yeast, COPII vesicles can transport transmembrane proteins from the ER to autophagic membranes (Shima et al., 2019). This raises the possibility that COPII-mediated transport similarly contributes to the delivery of SR transmembrane proteins to autophagosomes in DIOMs. However, if COPII-mediated vesicular transport were the predominant mechanism, blocking autophagy might be expected to result in the accumulation of vesicles positive for SR markers. In contrast, loss of autophagy caused extensive SR stacking. Thus, although COPII-mediated trafficking may contribute to the delivery of SR components, vesicular transport alone is unlikely to account for their transfer to autophagosomes.

Atg2-mediated lipid transfer is also unlikely to directly account for the transfer of SR transmembrane proteins. Atg2 is a bridge-like lipid-transfer protein that connects the ER and the growing phagophore, mediating bulk phospholipid transfer between these membranes (Osawa et al., 2022). Its elongated hydrophobic cavity provides a conduit for phospholipids but is not suited to transfer integral membrane proteins such as Sec61β or Cytob5. Thus, although Atg2 depletion causes prominent SR stacking, the phenotype is unlikely to result from the loss of direct transfer of SR transmembrane proteins by Atg2. Instead, impaired phagophore expansion upon Atg2 depletion may indirectly disrupt the transfer of SR membrane components to autophagic membranes through a distinct mechanism.

One attractive possibility is that SR components are transferred through transient membrane continuities between the SR and the phagophore. In mammalian cells, autophagosomes form in close association with PI3P-enriched ER subdomains called omegasomes (Axe et al., 2008; Nähse et al., 2024), and narrow tubular connections between the ER/omegasome and the growing phagophore have been observed (Hayashi-Nishino et al., 2009; Uemura et al., 2014; Ylä-Anttila et al., 2009). Such membrane continuities could allow integral membrane proteins to move laterally from the ER into autophagic membranes. Although an ortholog of the mammalian omegasome marker DFCP1 has not been identified in *Drosophila*, expression of human GFP-DFCP1 in the *Drosophila* fat body labels ring-shaped structures associated with Atg5- and Atg8-positive autophagic structures, suggesting that analogous membrane domains may also form in flies (Melani et al., 2017). We therefore speculate that transient tubular connections between the SR and the phagophore may allow SR lipids and transmembrane proteins to enter autophagosomal membranes during DIOM remodeling. Three-dimensional ultrastructural analyses, such as serial-section SEM and FIB-SEM, will be required to determine whether membrane continuities exist in DIOMs. Identifying the machinery responsible for forming the tubular connections will be important for testing their functional significance and for distinguishing direct membrane transfer from vesicular transport and Atg2- mediated lipid transfer.

Our genetic analyses further suggest that SR remodeling is closely coupled to early autophagic membrane formation and cannot be explained solely by receptor-mediated ER/SR- phagy. Disruption of core autophagy factors involved in initiating and expanding autophagic membranes, including FIP200, Atg1, Atg9, Atg2, and Atg18a, caused pronounced SR stacking, whereas disruption of the ubiquitin-like conjugation machinery, including Atg5, Atg7, and Atg8, resulted in substantially milder phenotypes (Fig. 2–3). Notably, the Atg8a^LDS^ mutant, which interferes with LIR-dependent cargo recognition, caused little SR stacking (Fig. 5D), arguing against canonical LIR–LDS-dependent selective autophagy as the primary mechanism underlying SR remodeling. Moreover, the accumulation of vesicular structures positive for SR markers together with autophagic membranes in Atg5-deficient cells suggests that at least part of the transfer of SR membrane components can proceed independently of the ubiquitin-like conjugation machinery. Together, these findings suggest the extent of SR stacking is associated with the impaired transfer of SR membrane components to nascent autophagic membranes (Fig. 5E).

This model is conceptually distinct from canonical receptor-mediated ER-phagy, in which portions of the ER are selectively recognized and sequestered within autophagosomes as cargo. Our observations suggest that, during DIOM remodeling, at least a fraction of the SR is instead incorporated into autophagosomal membranes themselves. These mechanisms are not necessarily mutually exclusive, and SR components may be eliminated both as cargo and as constituents of autophagic membranes. Autophagy may therefore remodel different organelles in DIOMs through distinct but coordinated mechanisms. While mitochondria are a major autophagic cargo during DIOM remodeling, the SR may simultaneously undergo autophagic turnover and provide membrane components for autophagosome formation. Autophagic membranes containing SR-derived components subsequently fuse with lysosomes and transform into the tubular autolysosomal network that forms in DIOMs (Fig. 5E) (Murakawa et al., 2020). As this network is eventually disassembled, SR-derived transmembrane proteins may be degraded through the ESCRT-dependent pathway or recycled. Because the SR is one of the most extensive membrane systems in muscle cells, its contribution to autophagic membranes may provide an efficient way to simultaneously remove excess SR membrane and support the high autophagic activity required for developmentally programmed muscle remodeling.

The requirement for autophagy in maintaining SR architecture appears to be evolutionarily conserved. As in *Drosophila* DIOMs, SR abnormalities are more pronounced upon disruption of early autophagy factors FIP200 or ULK1/ULK2 than upon disruption of ATG5 or ATG7 (Li et al., 2021; Masiero et al., 2009; Raben et al., 2008; Wang et al., 2019), suggesting a conserved relationship between early autophagic membrane biogenesis and SR homeostasis. However, determining whether mammalian SR membranes similarly contribute to autophagosome formation is technically challenging because mammalian cells possess partially redundant mechanisms for autophagosome–lysosome fusion (Matsui et al., 2018), multiple ER- phagy receptors (Chino and Mizushima, 2023; Mochida and Nakatogawa, 2022), and several Atg8-family paralogs (Schaaf et al., 2016). Future studies will therefore require approaches that can distinguish the incorporation of SR membrane components into autophagic membranes from their degradation through receptor-mediated SR-phagy.

Together, our findings reveal an unexpected link between SR remodeling and autophagosome biogenesis. Repurposing organelle membranes for autophagosome formation may provide an efficient mechanism to couple organelle turnover with large-scale membrane reorganization during developmentally programmed cellular remodeling.

## Materials and methods

### Reagents and antibodies

The following antibodies were used: mouse monoclonal anti-Dlg1 (1:100, DSHB, clone 4F3, RRID: AB_528203), rat monoclonal anti-GFP (1:200; GF090R, Nacalai Tesque, Kyoto, Japan), anti-mouse IgG Alexa Fluor 594 conjugate (1:400; A11005, Thermo Fisher Scientific, Waltham, MA), 1.4-nm Nanogold-conjugated goat anti-rat IgG Fab’ fragment (1:200; 2008-0.5 mL, Nanoprobes), and Alexa Fluor 546 Phalloidin (1:500; A22283, Thermo Fisher Scientific, Waltham, MA).

### Generation of transgenic flies

To generate UAS-GFP::Sec61β transgenic flies, cDNA encoding Sec61β and EGFP was amplified using the following primer sets; Fw-GFP-linker-Sec61β: 5’- GGATGAACTATACAAAGAATTCATGCCCGCTCCAGCCAGTTCAACG-3’, Rv-Sec61β - stop-vector: 5’-CCTTCACAAAGATCCTCTAGATTAAGAACGATTGTATTTGCCCCAAATG-3’ for Sec61β, and Fw-vec-kozak-GFP: 5’- TTCAGGCGGCCGCGGCTCGAGCACCATGAGTAAAGGAGAAGAACTTTTCAC-3’, Rv-GFP-stop-linker: 5’-GAATTCTTTGTATAGTTCATCCATGCCATGTG-3’ for EGFP. PCR amplification was performed using the DNA polymerase KOD One (TOYOBO, Osaka, Japan). The amplified fragments were assembled into linearized pJFRC7-20XUAS using *Xho*I and *Xba*I via Gibson Assembly. The resulting construct, pJFRC7-20xUAS-GFP-linker-Sec61β, was injected into embryos for phiC31-mediated insertion (Wellgenetics, Taiwan, Taipei).

### *Drosophila* strains and maintenance

Flies were reared under standard conditions at 25°C. For muscle-targeted gene expression, Mef2-GAL4 was used. The *w^1118^* strain was used as the wild-type control for the *ATG* mutants. UAS-*LacZ* was used as a control for UAS transgenes. All genetic combinations were generated by standard crosses. Detailed genotypes are described in Table S1. Genotypes of flies used in this study include the following:

1. *w^1118^*, referred to as wild-type (WT);
2. *Atg5^5cc5^*/*FM7_GFP* (from J. H. Lee) (Kim et al., 2016), referred to as *Atg5* null;
3. *Atg9^d51^*/*CyO_GFP* (from G. C. Chen) (Wen et al., 2017), referred to as *Atg9* null;
4. *Atg9^Gal4KO^*/*CyO_GFP* (from G. C. Chen) (Wen et al., 2017), referred to as *Atg9* null;
5. *FIP200^3F5^*/*TM6B_GFP* (from J.H. Lee) (Kim et al., 2013), referred to as *FIP200* null;
6. *FIP200^4G7^*/*TM6B_GFP* (from J.H. Lee) (Kim et al., 2013), referred to as *FIP200* null;
7. *Atg8^Δ12^*/*FM7_GFP* (from G. Juhasz and A. Szabo) (Vincze et al., 2026), referred to as *Atg8a* null;
8. *Atg8a^K48A,Y49A^* (from I. P. Nezis) (Rahman et al., 2022), referred to as *Atg8a^LDS^*;
9. *y,w*; *P*{*w^+mC^=GAL4-Mef2.R*}3 (BDSC_27390), referred to as Mef2-GAL4;
10. *w*; *P*{*w^+mC^=UAS-lacZ.B*}*Bg4-1-2* (BDSC_1776), referred to as LacZ;
11. *w*; *P{w[+mC]=UAS-Dcr-2.D}10* (BDSC_24651), referred to as UAS-Dcr2;
12. *w*; *P*{*w*[*+mC*]*=UAS-mCD8::GFP.L*}*LL5, P*{*UAS-mCD8::GFP.L*}*2* (BDSC_5137), referred to as mCD8-GFP;
13. *w*; *P{w+=UAS-mCherry::PH^PLCδ^}* (from A. A. Kiger), referred to as mCh-PH^PLCδ^;
14. *w*; *P{y[+7.7] w[+mC]=20XUAS-tdTomato::Sec61β}attP2* (BDSC_64747), referred to as tdTomato-Sec61β;
15. *w*; *P{w[+mC]=UAS-GFP::ER-SR}18* (BDSC_59041), referred to as GFP-Cytob5;
16. *w*; *UAS-GFP::Sec61β^attP40^* (this study), referred to as GFP-Sec61β;
17. *w*; *P{w[+mC]=UASp-Ggal/LYZ.GFP.KDEL}3/TM6B, Tb[1]*, referred to as Lyz-GFP- KDEL;
18. *y,w*; *UAS-GFP::Atg8a* (from T. Neufeld), referred to as GFP-Atg8a;
19. *w*; *UASt-GFP::Stx17^4^* (Fujita et al., 2017), referred to as GFP-Stx17;
20. *y,v*; *P*{*y*[*+t7.7*] *v*[*+t1.8*]*=TRiP.HMS01611*}*attP2*/*TM3, Sb*[*1*] (TRiP, BDSC_36918), referred to as FIP200 RNAi;
21. *w*; *UAS-IR-Atg2^KK100148^* (VDRC_108447), referred to as Atg2 RNAi;
22. *w*; *UAS-IR-Atg18a^KK100064^* (VDRC_105366), referred to as Atg18a RNAi;
23. *w*; *UAS-IR-Atg1* (from G. C. Chen), referred to as Atg1 RNAi;
24. *y,v*; *P*{*y*[*+t7.7*] *v*[*+t1.8*]*=TRiP.HMS01246*}*attP2* (BDSC_34901), referred to as Atg9 RNAi;
25. *y,v*; *P*{*y*[*+t7.7*] *v*[*+t1.8*]*=TRiP.HMS01358*}*attP2*/*TM3, Sb*[*1*] (BDSC_34369), referred to as Atg7 RNAi;
26. *w*; *UAS-IR-Atg5* (NIG-Fly_1643R-2), referred to as Atg5 RNAi;
27. *y,v*; *P*{*y*[*+t7.7*] *v*[*+t1.8*]*=TRiP.JF01937*}*attP2* (BDSC_25896), referred to as Stx17 RNAi;
28. *y,v*; *P*{*y*[*+t7.7*] *v*[*+t1.8*]*=TRiP.JF01883*}*attP2* (BDSC_25862), referred to as SNAP29 RNAi;

### Immunofluorescence staining

Muscle preparations were performed as previously described (Ribeiro et al., 2011). 3IL or staged pupae were pinned flat on Sylgard-coated Petri dishes in dissection buffer (5 mM HEPES, 128 mM NaCl, 2 mM KCl, 4 mM MgCl_2_, 36 mM sucrose; pH 7.2). The dissected preparations were fixed in 4% paraformaldehyde (PFA) in phosphate buffer (Nacalai Tesque, Kyoto, Japan, #09154-85) for 20 min at room temperature. After fixation, the samples were unpinned and blocked in PBS containing 0.3% bovine serum albumin and 0.6% Triton X-100 for 5 min at room temperature. Samples were incubated with the anti-Dlg1 primary antibody overnight at room temperature, washed with PBS, and incubated with the Alexa Fluor-conjugated secondary antibody (Thermo Fisher Scientific, Waltham, MA) for 2 h at room temperature. The samples were then washed with PBS and mounted in FluorSave reagent (Merck Millipore, Darmstadt, Germany).

### Confocal fluorescence microscopy

For imaging of pupal DIOMs, staged pupae were removed from their pupal cases and mounted between a glass slide and a coverslip (Murakawa et al., 2020). DIOMs were imaged through the dorsal cuticle. Both live and fixed samples were imaged using an FV3000 confocal microscope (EVIDENT, Tokyo, Japan) equipped with a 60× silicone/1.30 NA UPlanSApo silicone-immersion objective (EVIDENT, Tokyo, Japan). Images were acquired using Fluoview software (EVIDENT, Tokyo, Japan) and processed and analyzed using ImageJ (NIH, Bethesda, MD, USA).

### TEM

Staged pupae were removed from their pupal cases, pinned on a Sylgard-covered Petri dish, and dissected in fixative containing 2% PFA, 2.5% glutaraldehyde, 150 mM sodium cacodylate, 5 mM calcium chloride (pH 7.4). The dissected samples were fixed for 2 h at room temperature and then overnight at 4°C. Samples were washed with 0.1 M phosphate buffer (PB; pH 7.4), post-fixed in 1% OsO_4_ in 0.1 M PB for 2 h, dehydrated through a graded ethanol series, and embedded flat in Epon 812 (TAAB, Aldermaston, UK). Ultrathin sections (70 nm) were collected on Formvar-coated copper grids (Nisshin EM, Tokyo, Japan), double-stained with uranyl acetate and lead citrate, and then observed using a JEM-1400Flash transmission electron microscope (JEOL, Tokyo, Japan).

### Immuno-EM

For immunogold labeling, staged pupae were rapidly frozen in liquid nitrogen and cryosectioned using a cryostat. The sections were fixed with 4% PFA in 0.1 M phosphate buffer (PB). After permeabilization and blocking with a solution containing 50 mM NH₄Cl, 0.1% saponin, and 1% BSA in PB, the sections were incubated with an anti-GFP antibody (1:200; GF090R, Nacalai Tesque), followed by a 1.4-nm Nanogold-conjugated goat anti-rat IgG Fab′ fragment (1:200; 2008, Nanoprobes). Silver enhancement was performed according to a pre-embedding immunogold labeling protocol as previously described (Yoshimori et al., 2000).

### Image analyses

For quantification of the size of SR-marker-positive objects (Fig. 2B′, 2C′, and 5D′), the entire DIOM was manually outlined using the Freehand Selection tool in ImageJ and defined as the region of interest (ROI). The GFP channel was filtered and background-subtracted, followed by thresholding to identify GFP-positive objects. The area of individual GFP-positive objects within the ROI was measured using the Analyze Particles function in ImageJ. The mean object area (µm²) was calculated for each DIOM and used for statistical analysis. At least 15 DIOMs from five individual animals were analyzed for each genotype.

For 3D reconstruction (Fig. 1—figure supplement 1B), the GFP channel was filtered and background-subtracted, followed by thresholding to generate binary images of GFP-positive objects. The processed z-stack images were then surface rendered using the 3D Viewer in ImageJ.

For quantification of vesicle density (Fig. 4C′), the entire DIOM was manually outlined using the Freehand Selection tool in ImageJ and defined as the ROI. After background subtraction, the DIOM area was measured using the Measure function. Images were then processed using the Convolve function (Process > Filters > Convolve) and converted to binary images. Vesicular structures with diameters ranging from 0.732 to 1.218 µm were detected by Hough circle transformation using the Hough Circle Transformation plugin (Plugins > Hough Circle Transformation). The number of detected vesicles was normalized to the DIOM area and expressed as the number of vesicles per unit area. 15 DIOMs from five individual animals were analyzed for each genotype.

## Statistics

Statistical analyses were performed using GraphPad Prism version 10.5.0 (GraphPad software, Boston, MA). Data are presented as means unless otherwise indicated. Comparisons between two groups were performed using the Mann–Whitney U test. For comparisons involving more than two groups, one-way analysis of variance (ANOVA) followed by Dunnett’s multiple comparisons test was used. P values < 0.05 were regarded as statistically significant. All experiments were performed in parallel with the corresponding controls.

## Acknowledgments

We are grateful to Jun H. Lee (University of Michigan); Gábor Juhász and Áron Szabó (HUN-REN Biological Research Centre); Guang-Chao Chen (Academia Sinica); Ioannis P. Nezis (University of Warwick); the Bloomington Drosophila Stock Center; the Vienna Drosophila Resource Center (VDRC); the Drosophila Genomics Resource Center (DGRC); and NIG-FLY for providing reagents. We thank Amy Kiger (University of California, San Diego) for helpful discussions; Hiroki Takechi (Institute of Science Tokyo) for technical assistance; the Biomaterials Analysis Division of the Institute of Science Tokyo for DNA sequencing; and Marija Landekic (McGill University) for English editing. This work was supported in part by JSPS KAKENHI grants 21H05147 and 26H01620 (to N.F.), AMED-PRIME 24gm6410016h0001 (to N.F.), and a research grant from the Takeda Science Foundation (to N.F.).

## Author contributions

K.: Investigation, Formal analysis, Data curation, Methodology, Writing – review and editing.
Borthakur: Investigation, Writing – review and editing.
Bennani: Investigation, Formal analysis, Data curation, Writing – review and editing.
M.: Methodology, Writing – review and editing.
S.: Methodology, Investigation, Writing – review and editing.
F.: Conceptualization, Methodology, Investigation, Formal analysis, Supervision, Funding acquisition, Project administration, Writing – original draft, Writing – review and editing.

## Supplementary figure legends

Figure 1—figure supplement 1

**Time-lapse imaging of GFP-Cytob5 in DIOMs during the early pupal stage**

A. Time-lapse imaging of GFP-Cytob5 in control and *Atg2* RNAi DIOMs between 13 and 22 h APF. (B) 3D reconstruction of live GFP-Cytob5 imaging in *Atg2* RNAi DIOMs at 22 h APF. Images were acquired by confocal microscopy at 90-s intervals and reconstructed in 3D using ImageJ.

Figure 3—figure supplement 1 TEM of DIOMs

A. TEM image of a longitudinal section of *FIP200*-null DIOM at 22 h APF. Stacked SR in pink. (B) TEM images of transverse sections of WT and *Atg5*-null DIOMs at 4 d APF. Myofibrils in green.

Figure 5—figure supplement 1

**Colocalization of tdTomato-Sec61β with GFP-Stx17 upon SNAP29 RNAi** Colocalization of tdTomato-Sec61β with GFP-Stx17 in *SNAP29* RNAi DIOMs at 4 d APF. Line profiles along the white dotted arrow are shown.

Supplementary Table 1: Detailed *Drosophila* genotypes used in each figure

Supplementary File 1: Raw data files corresponding to all quantified results presented in the manuscript.

## Abbreviations used

APF: after puparium formation;
ATG: autophagy-related
COPII: coat protein complex II
DIOM: dorsal internal oblique muscle
LBWM: larval body wall muscle
LDS: LIR-docking site
LIR: LC3-interacting region
mCh: monomeric Cherry
SR: sarcoplasmic reticulum
TEM: transmission electron microscopy
WT: wild-type
3IL: third- instar larval stage.

