## Supplementary Figures for "Sarcoplasmic reticulum remodeling through autophagosome biogenesis during *Drosophila* metamorphosis"

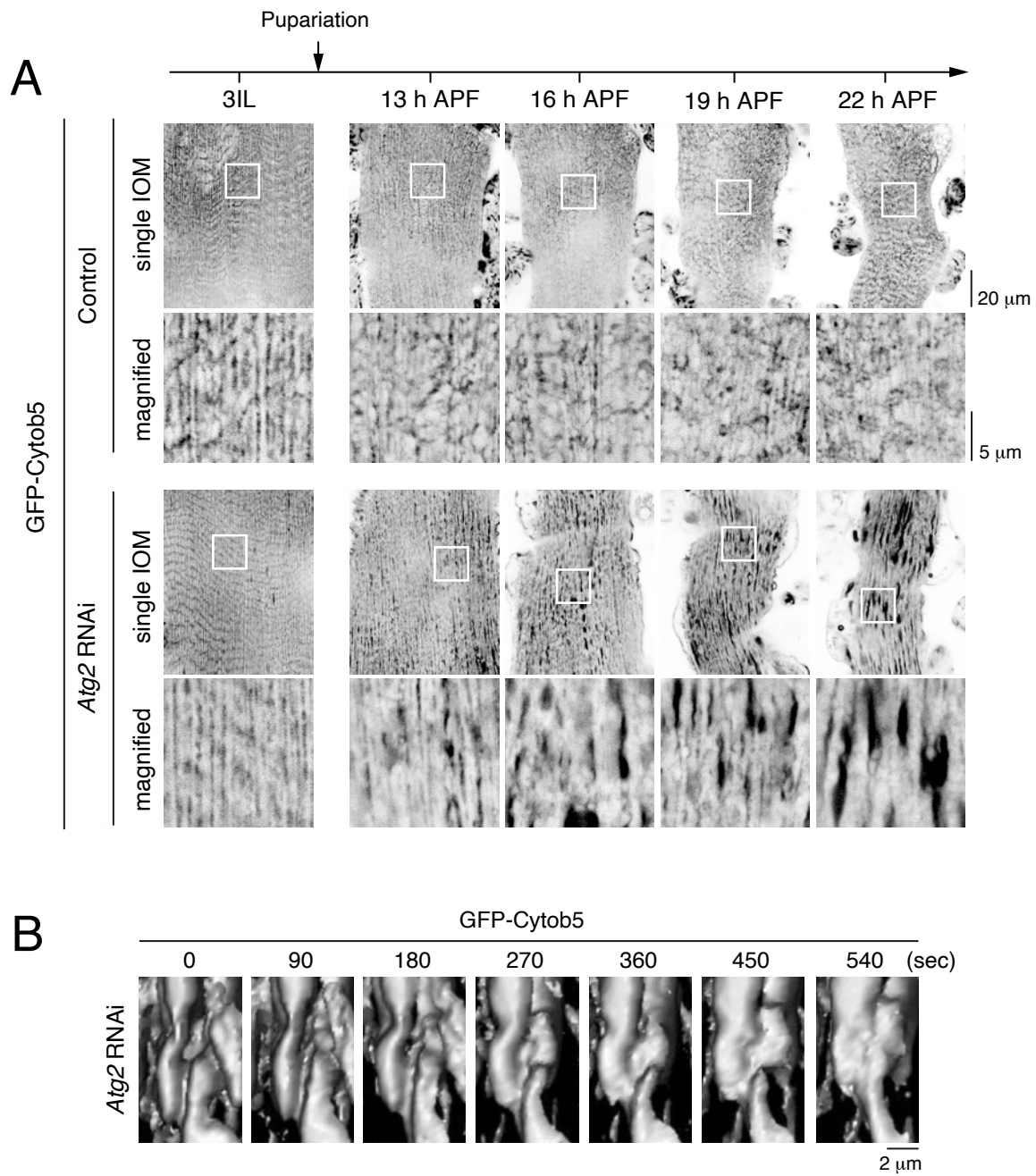

**Figure 1—figure supplement 1**

**Time-lapse imaging of GFP-Cytob5 in DIOMs during the early pupal stage**

(A) Time-lapse imaging of GFP-Cytob5 in control and *Atg2 RNAi* DIOMs between 13 and 22 h APF. (B) 3D reconstruction of live GFP-Cytob5 imaging in *Atg2 RNAi* DIOMs at 22 h APF. Images were acquired by confocal microscopy at 90-s intervals and reconstructed in 3D using ImageJ.

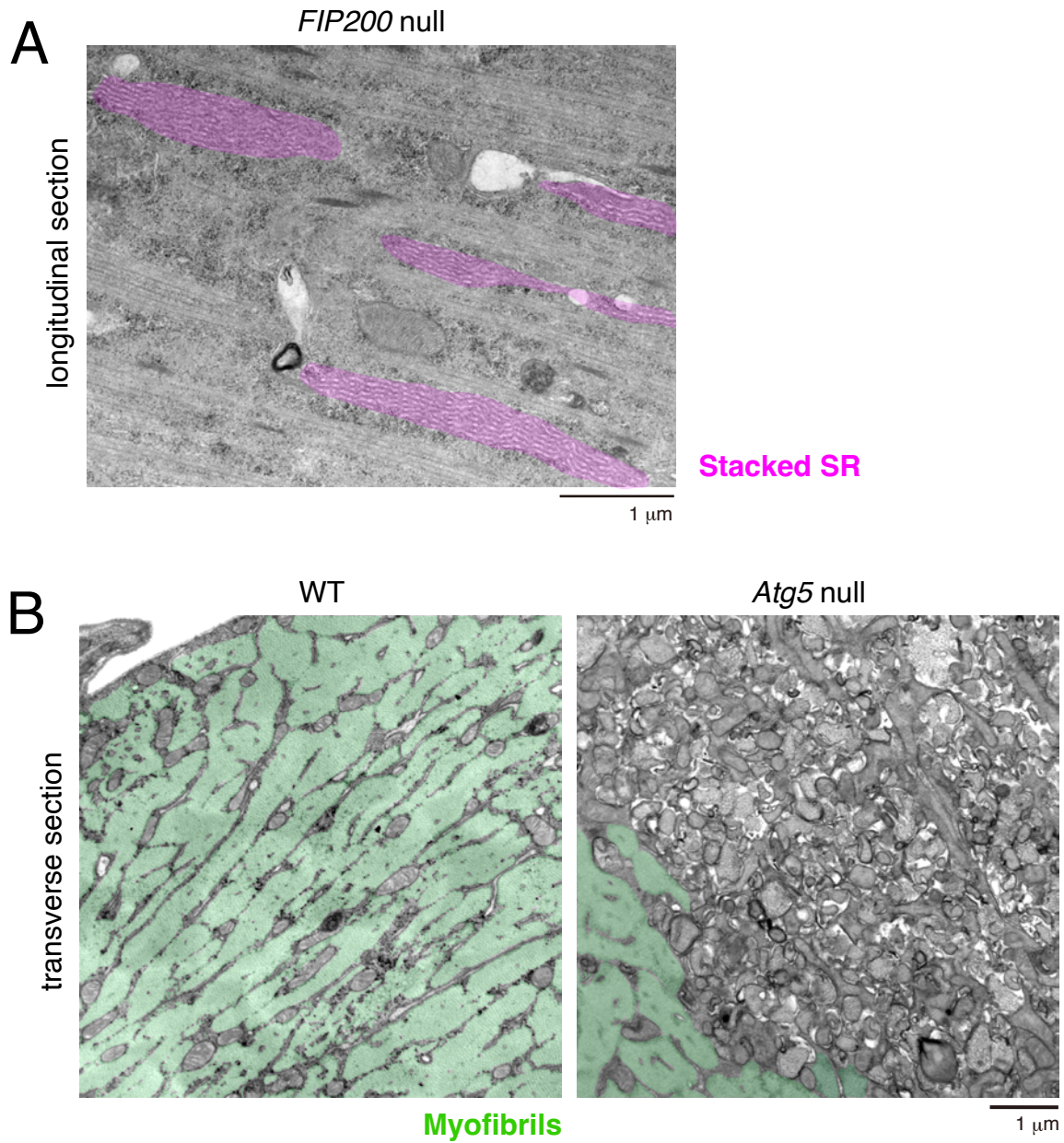

**Figure 3—figure supplement 1**

**TEM of DIOMs**

(A) TEM image of a longitudinal section of *FIP200*-null DIOM at 22 h APF. Stacked SR in pink. (B) TEM images of transverse sections of WT and *Atg5*-null DIOMs at 4 d APF. Myofibrils in green.

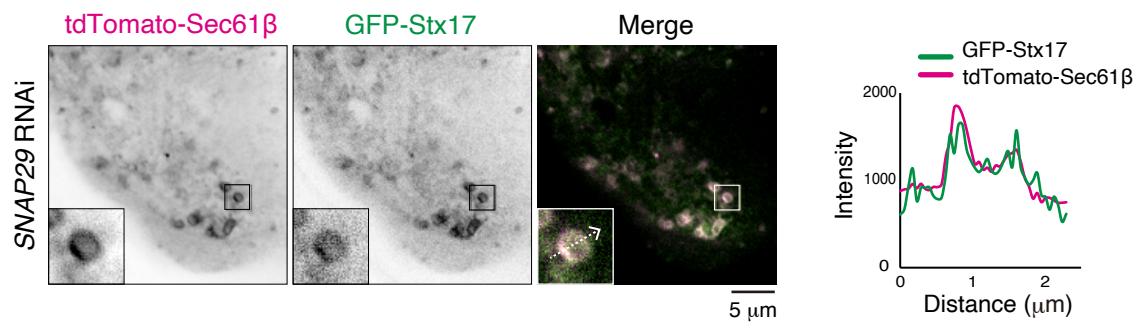

### Figure 5—figure supplement 1

#### Colocalization of tdTomato-Sec61β with GFP-Stx17 upon SNAP29 RNAi

Colocalization of tdTomato-Sec61β with GFP-Stx17 in *SNAP29* RNAi DIOMs at 4 d APF. Line profiles along the white dotted arrow are shown.
